# Not Quite All in Our Head: Intervention is a Better Predictor of tDCS Mind-Wandering Effects than Subjective Beliefs About Experimental Results

**DOI:** 10.1101/2021.11.14.468553

**Authors:** Matilda S. Gordon, Paul E. Dux, Hannah L. Filmer

## Abstract

**Background:** Establishing adequate blinding for non-invasive brain stimulation research is a topic of extensive debate, especially regarding the efficacy of sham control methods for transcranial direct current stimulation (tDCS) studies. Fassi and Cohen Kadosh [1] assessed the influence of subjective participant belief regarding stimulation type (active or sham) and dosage on behaviour using data from Filmer et al. [2] who applied five stimulation protocols (anodal 1.0mA, cathodal 1.0mA, cathodal 1.5mA, cathodal 2.0mA and sham) to assess the neural substrates of mind wandering. Fassi and Cohen Kadosh [1] concluded that subjective belief drove the pattern of results observed by Filmer et al. [2].

**Objective:** Fassi and Cohen Kadosh [1] did not assess the key contrast between conditions in Filmer et al. (2019) – 2mA vs sham – rather they examined all stimulation conditions. Here, we consider the relationship between objective and subjective intervention in this key contrast.

**Methods:** We replicated the analysis and findings of both Filmer et al. [2] and Fassi and Cohen Kadosh [1] before assessing 2mA vs. sham via Bayesian ANOVA on subjective belief regarding stimulation type and dosage.

**Results:** Our results support objective intervention as the strongest predictor of stimulation effects on mind-wandering when 2mA vs sham was examined, over and above that of subjective intervention.

**Conclusions:** The conclusions made by Filmer et al. [2] are confirmed. However, it is important to control for and understand the possible effects of subjective beliefs in sham-controlled studies. Best practice to prevent these issues remains the inclusion of active control conditions.

---

Non-invasive brain stimulation (NIBS) is a popular tool for investigating causal relationships between activity in cortical brain regions and behaviour [4]. There are multiple methods of NIBS, one of the most common being transcranial direct current stimulation (tDCS [5]). Interest has grown in this brain stimulation approach as it can lead to cognitive enhancement both within clinical settings, such as for the treatment of drug-resistant depression [6] [7], as well as in commercial settings via do-it-yourself devices such as the *foc.us* headset [8] [9] [10]. In 2020 the brain stimulation industry was projected to be worth an estimated three billion dollars [11], a figure that is almost certain to increase in the future. Given this broad interest there is growing demand for clarity on the efficacy of brain stimulation across the extensive stimulation parameter space.

tDCS typically involves passing electrical current through two electrodes; a cathode and an anode [2] allowing for causal inferences to be made regarding the stimulated brain region(s) and behavioural performance on a task(s). A relatively cost-effective method both in terms of temporal and financial considerations, tDCS has demonstrated promising effects on a range of functions such as motor [12] and speech motor learning [13]. In addition, a variety of cognitive operations that have previously been shown to be influenced by tDCS include working memory [14] [15] response selection [16], multitasking [17] and attention [18] [19]. Given its flexibility and effects on a wide range of brain regions and behavioural processes, tDCS shows promise in both pure research and applied settings.

Although tDCS has been employed extensively, some reviews and meta-analyses [20] [21] [22] have suggested limited to no effects of transcranial stimulation, with a key focus being on large variability across participants. For example, Lopez-Alonso et al., [21] found only 45% of participants (n=56) responded to anodal transcranial current stimulation as expected. In an opinion piece, Filmer et al., [5] provide recommendations to protect the reliability, reproducibility, and validity of effects of NIBS studies, whether they yield significant results or not. Key factors identified by the authors to be controlled include poor methodological design, under-powered samples which give rise to inflated results and, most importantly for the present study, the inadequate blinding of control conditions.

The most common method of blinding in tDCS studies is the sham control method [23], which mimics the typical initial sensations (i.e., itching or tingling) induced by tDCS by delivering active stimulation for a short period at the beginning (and sometimes again at the end), then either no stimulation or very reduced pulses (which allow some continued sensation) for the rest of the session. Typically, the length of this period of active stimulation is dependent on the stimulation length used for the active condition [24]. It is assumed that sham stimulation controls for any potential unrelated effects of the direct cortical stimulation [24]. However, the efficacy of sham-controlled approaches has been called in to question. Indeed, in NIBS studies, participants can report perceptual sensations such as visual disturbances, and cutaneous feelings [25] [26]. Cutaneous feelings are common in tDCS as seen in a prospective comparison conducted by Kessler et al. [26] with 131 subjects in 277 tDCS sessions. Such feelings were significantly higher in active conditions as compared to sham, for example tingling (89% active vs. 53% sham) and itching (81% vs. 42%). This is potentially a major issue for sham-controlled studies, as observed results may reflect peripheral effects rather than the influence of stimulation on the cortex. Or, put differently, failure of the blinding condition (although, see [27] and [28]). Another factor that may add to blinding inefficiency is the inadequate reporting of adverse events (such as cutaneous feelings), as this is not only a safety concern but also prevents the experimenter from gaining an understanding of the strength of blinding [25]. Indeed, Wallace et al. [29] found that when investigating the comfort and efficacy of sham-blinding, after a second session of tDCS participants were able to correctly guess stimulation above chance (65%).

Given these issues of blinding, Fassi and Cohen Kadosh [1] suggested that participants’ subjective feeling regarding the stimulation they received may alter their task performance. Specifically, these authors investigated measures that demonstrate subjective opinion of both stimulation mode (active or sham) and dosage level (not applicable, low, moderate, or high). This was done as the standard practice in the field to evaluate blinding is to analyse whether participants guesses regarding their allocated condition (active vs. sham or dosage level) relative to chance and then analyse behavioural data independently of this measure. Including measures of blinding success within the main analysis of behavioural data, whether this be via a covariate or a similar method, may be a more sensible and sensitive approach to investigating blinding efficacy.

Fassi and Cohen-Kadosh [1] used open-access data from our group, Filmer et al. [2], which involved the sustained attention to response task (SART) along with mind-wandering probes. The original study [2] investigated the effect of tDCS intensity and polarity on mindwandering. Five variations of polarity and intensity, including a sham condition were randomly assigned to participants in a between-subjects design. The effect of stimulation intensity (objective intervention) and stimulation dosage (objective dosage) was investigated using Bayesian statistics and a linear relationship was found between stimulation dosage and mind wandering as assessed via the propensity for participants to report task unrelated thoughts (TUT). Of particular importance for the present study, only the strongest stimulation intensity, cathodal 2.0mA stimulation, had a reliable effect with moderate evidence for stimulation increasing mind-wandering.

Fassi and Cohen-Kadosh [1] sought to investigate the hypothesis that “subjective intervention” (participants’ subjective beliefs about the intervention they received) and “subjective dosage” (participants subjective beliefs about the stimulation intensity they received) drove the effect we previously observed. Using a Bayesian ANOVA to compare objective intervention and subjective intervention for average mind-wandering, Fassi and Cohen-Kadosh [1] concluded that subjective participant belief regarding the type of stimulation received (sham versus active) was a better predictor of participant performance than objective intervention, subjective and objective intervention combined or an interaction between the two. Particularly, those who believed they had received active stimulation had higher levels of mind wandering than those who answered sham. Regarding dosage, a Bayesian ANOVA comparing objective intervention with subjective dosage revealed participant’s subjective dosage beliefs were better at explaining mind wandering scores than objective intervention, subjective intervention and objective intervention combined or an interaction between the two. With further investigation via a Bayesian ANOVA with only subjective dosage, it was argued that as subjective dosage increased, so did average mindwandering score in a proportional manner.

On the surface, the results of Fassi and Cohen-Kadosh [1] call into question those of Filmer et al. [2] and, more broadly, those of any study with purely a sham control. However, a key limitation of the Fassi and Cohen-Kadosh [1] approach was that they examined all conditions on the Filmer et al. [2] study in a single analysis, whereas only effects were observed for cathodal 2.0mA stimulation as compared to sham comparison with moderate evidence (see Table 1 for specific BF_10_ values for the experimental conditions from Filmer et al. [2]). The present investigation replicated the analyses of Fassi and Cohen Kadosh [1] however focussed at this key effect.

**Table 1.** Individual BF_10_ and percent error for each stimulation condition (as compared to sham) for average task-unrelated thought across all experimental trials [2].

| **Stimulation Condition** | **$BF_{10}$** | **Error %** |
| --- | --- | --- |
| Anodal 1.0mA | 0.781 | 0.010 |
| Cathodal 1.0mA | 0.540 | 0.007 |
| Cathodal 1.5mA | 2.189 | 0.005 |
| Cathodal 2.0mA | 7.436 | 8.694e-5 |

The present study predicted that once this issue was accounted for, the effects of subjective belief described by Fassi and Cohen Kadosh [1] would not hold. To preview the results, when evaluating only cathodal 2.0mA as compared to sham the effects described by Fassi and Cohen Kadosh [1] were not observed: objective intervention was the strongest predictor of mind-wandering within the model, over and above the other included factors. Similarly for dosage, objective dosage provided the best model fit over and above that of subjective dosage, a combination of objective and subjective dosage, or an interaction between the two.

## Method

### Data and Materials

The study by Filmer et al. [2] was pre-registered via the Open Science Framework with details of analysis plan, methodology and sample size by the authors (https://osf.io/j6mqa/). The raw data from the original study has been made open source by the authors and can be accessed via the UQ eSpace [3], as well as the demographic and questionnaire data used to establish the experimental measures. Experimental materials such as the task can also be accessed here. For a full overview of the original methods, refer to Filmer et al. [2].

### Subjects

One hundred and fifty subjects (mean age= 23, SD= 5, 96 females) participated in the study. All subjects were right-handed with normal or corrected to normal vision. Subjects were assigned to one of five different stimulation groups based on their participant number sequentially (subject 1, 6, 11 etc. were assigned to anodal stimulation). The final sample per group was 30 [2].

### Task (Filmer et al., 2019a [2])

Subjects completed a sustained attention-to-cue task (SART), responding via a keyboard key press (space bar) to non-target stimuli (any number except 3). In each trial, a stimulus was presented in the centre of the display. Subjects were to withhold their answer when the target stimulus was presented (the number 3). Stimuli were presented for 1s and a 1.2s blank screen appeared between stimuli. The background of the display was light grey (RGB: 104, 104, 104), the stimuli were black (RGB: 255, 255, 255), in size 40 font. Trials consisted of an average of 20 non-target stimuli (*SD*= 5.69). At the end of half of the trials a target stimulus was presented and in the other half an unrelated thought probe was presented in the centre of the display in size 20 black font. Task un-related thought (TUT) probes asked: *“To what extend have you experienced task unrelated thoughts prior to the thought probe? 1 (Minimal) – 4 (Maximal)”*. The corresponding numbers on a keyboard were used to indicate their response. Subjects undertook two practice trials prior to stimulation (one target and one thought probe). A total of 48 trials were completed after stimulation with 24 target and 24 thought probes, split into 8 blocks (6 trials each) with approximately three of each trial type randomly intermixed.

### tDCS

tDCS was administered using a Neuroconn stimulator via two 5 × 5cm saline-soaked electrodes. The reference electrode was located on the right orbito-frontal region and the target electrode was placed over F3 (EEG 10-20 system). The four active stimulation groups consisted of various polarity and intensity combinations: anodal 1.0mA, cathodal 1.0mA, cathodal 1.5mA, and cathodal 2.0mA. In these conditions, total stimulation duration was 20 minutes including a 30s ramping up and down period. Those in the sham stimulation condition also received a 30s ramping period, but only received 15s of active stimulation before stimulation ramped down for a further 30s (thus a total of 1 minute and 15 seconds of stimulation). During stimulation, subjects were asked to sit quietly with their eyes open.

### Statistical Analysis

As with both Filmer et al. [2] and Fassi and Cohen Kadosh [1] all statistical analysis was conducted in JASP (version 0.14.1 for MacOS [30]). Analyses were conducted on the open-access dataset from Filmer et al. [3] using average mind wandering scores calculated from the whole experimental session for each subject.

To verify both the data and the coding were reproducible, we implemented the statistical analyses of both Fassi and Cohen Kadosh [1] and Filmer et al. [2]. All previous findings were replicated. Bayesian statistics and their frequentist counterparts were used as both were included in the previous papers. BF_10_ ~1 values provide no evidential value, BF_10_ of 1-3 were interpreted as anecdotal, BF_10_ of 3-10 as moderate, BF_10_ >10 as strong evidence favouring the alternate hypothesis that the examined model provides a better fit than the null model. BF_01_ values provide support for the null hypothesis and thus BF_01_ 1-3 was interpreted as anecdotal, BF_01_ 3-10 as moderate, and lastly BF_01_ >10 as strong evidence. As frequentist statistics were also evaluated, all values of *p*< .05 were accepted as statistically significant.

In JASP (using default priors) we replicated the Bayesian ANOVAs conducted by Fassi and Cohen Kadosh [1] however, as previously outlined, only data from the cathodal 2.0mA and sham condition were included in our key analysis. All relevant post-hoc pairwise comparisons were conducted following each analysis. The first Bayesian ANOVA included objective intervention and subjective intervention as between-subjects factors. To rule out the possibility of subjective information influencing performance in the absence of an effect, this analysis was also performed for each individual condition with no significant effect compared to sham (anodal 1.0mA, cathodal 1.0mA and cathodal 1.5mA). The second Bayesian ANOVA employed objective intervention and subjective dosage as the between-subject factors. Although the authors conducted a third Bayesian ANOVA to investigate the effect of subjective dosage only, we did not include this given the results of the second ANOVA indicating subjective dosage has no evidential value of an effect on mind-wandering.

## Results

### Summary of the Results from Fassi and Cohen Kadosh [1]

The authors compared model fit via three Bayesian ANOVAs. The first ANOVA conducted included objective intervention and subjective intervention as between-subject factors. They posited subjective intervention (BF_10_= 3.374, *t*(148)= 2.55, *p*= .012) as being the best predictor of the observed changes in mind-wandering over and above that of objective intervention. It was concluded by the authors that subjective participant belief regarding the type of stimulation received (sham versus active) was a better explanation of participant performance than objective intervention, subjective and objective intervention combined or an interaction between the two. The full model result from this ANOVA can be found in the table located in Appendix A1. The second ANOVA featured objective intervention and subjective dosage as the between-subject factors, and average TUT across all trials as the outcome measure. The authors concluded that participant belief about stimulation dosage was a better predictor of mind wandering than objective intervention alone, a combination of the two measures or an interaction between them (BF_10_= 3.708). The authors did not report the frequentist statistics for this finding however when we replicated the ANOVA we determined the values to be BF_10_= 3.713, *F*(3, 146)= 3.829, *p*= .011. The full model table can be found in Appendix A2. When investigating dosage alone, subjective dosage (BF_10_= 5.911, *F*(3,198)= 4.198, *p*=.007) was also a better predictor of mindwandering changes over and above that of objective dosage. It was concluded by the authors via a linear regression (BF_10_= 24.26, β= .263, *t*(148)=3.13, *p*= .001) that as participants subjective belief about the dosage they received increased (i.e., sham, weak, moderate and then strong) mind-wandering increased proportionally, thus subjective dosage is a significant predictor of mind-wandering.

### Effect of Subjective Belief on Mind-Wandering via 2.0mA Cathodal Stimulation

We implemented the first Bayesian ANOVA discussed above, employing objective intervention and subjective intervention as between-subject factors and average TUT as the outcome measure, however, crucially, we excluded the conditions showing limited evidence of differences between stimulation and sham, evaluating only cathodal 2.0mA vs the sham control condition. The full model comparing against the null can be found in Appendix B1 however a summarised model table comparing these results to those of Fassi and Cohen Kadosh [1] can be found in Table 2. We observed that objective intervention was the strongest model predictor with moderate evidence (BF_10_= 7.436, *F*(1, 58)= 8.263, *p*=.006) when accounting only for the conditions that previously produced meaningful effects. Subjective intervention proved to be the least predictive within this model (BF_10_= 0.276, *F*(1, 58)= 0.104, *p*= .748). Post-hoc pairwise comparisons on the objective information indicated that cathodal 2.0mA had increased mind-wandering (*M*= 2.288, BF_10_= 7.436, *t*(59)= −2.866, *p*= .006) as compared to sham (*M*= 1.772).

**Table 2.**
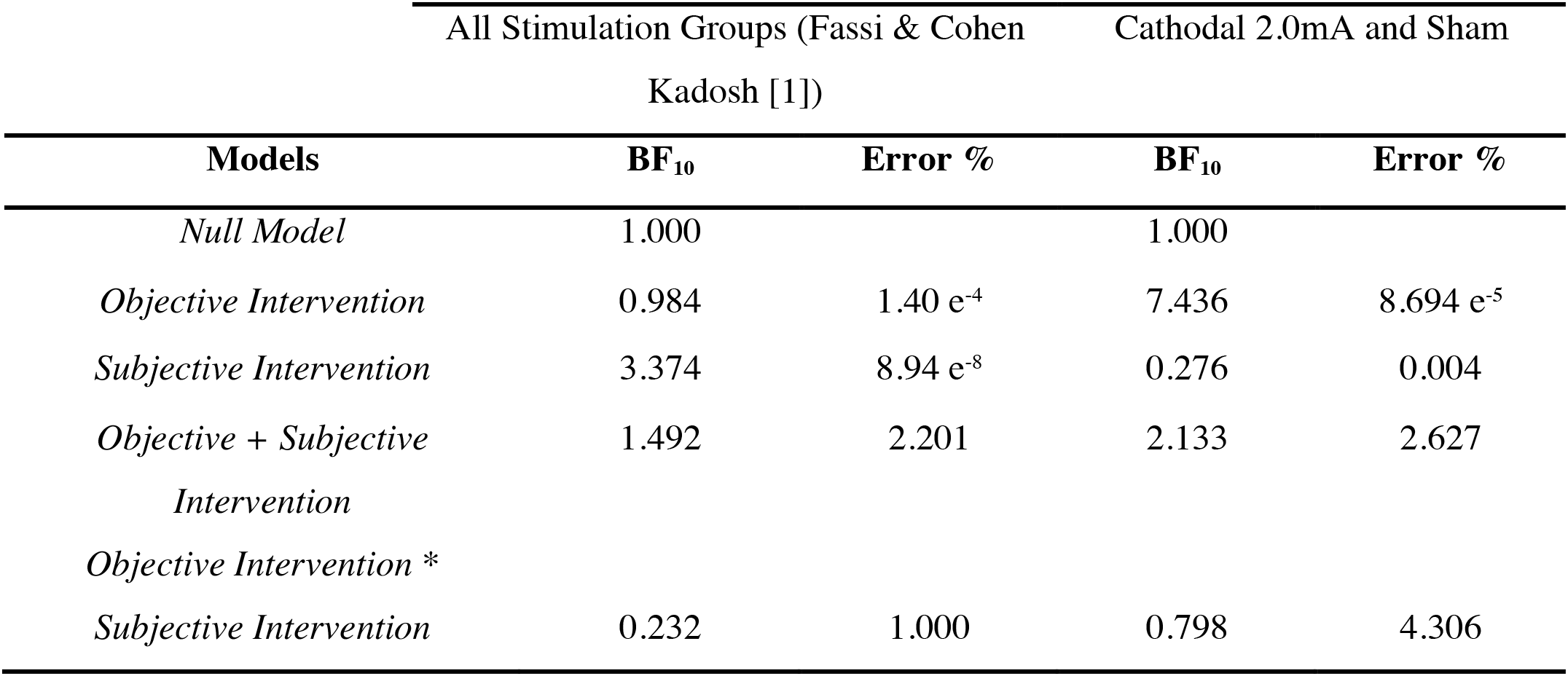
Comparison of the summarised Bayesian ANOVA of objective intervention and subjective intervention findings from Fassi & Cohen Kadosh [1] and the present study evaluating cathodal 2.0mA stimulation only.

When comparing objective intervention to subjective dosage, again objective intervention was the strongest predictor of mind-wandering within the model (BF_10_= 7.436, *F*(1, 58)= 8.263, *p*=.006). Subjective dosage was the weakest predictor in the model (BF_10_= 0.254, *F*(3, 56)= 0.847, *p*=.474). A comparison of these model findings compared to those found by Fassi and Cohen Kadosh [1] can be found in Table 3, which demonstrates that when evaluating only conditions previously found to demonstrate meaningful results, the effect of subjective dosage can no longer be observed.

**Table 3.**
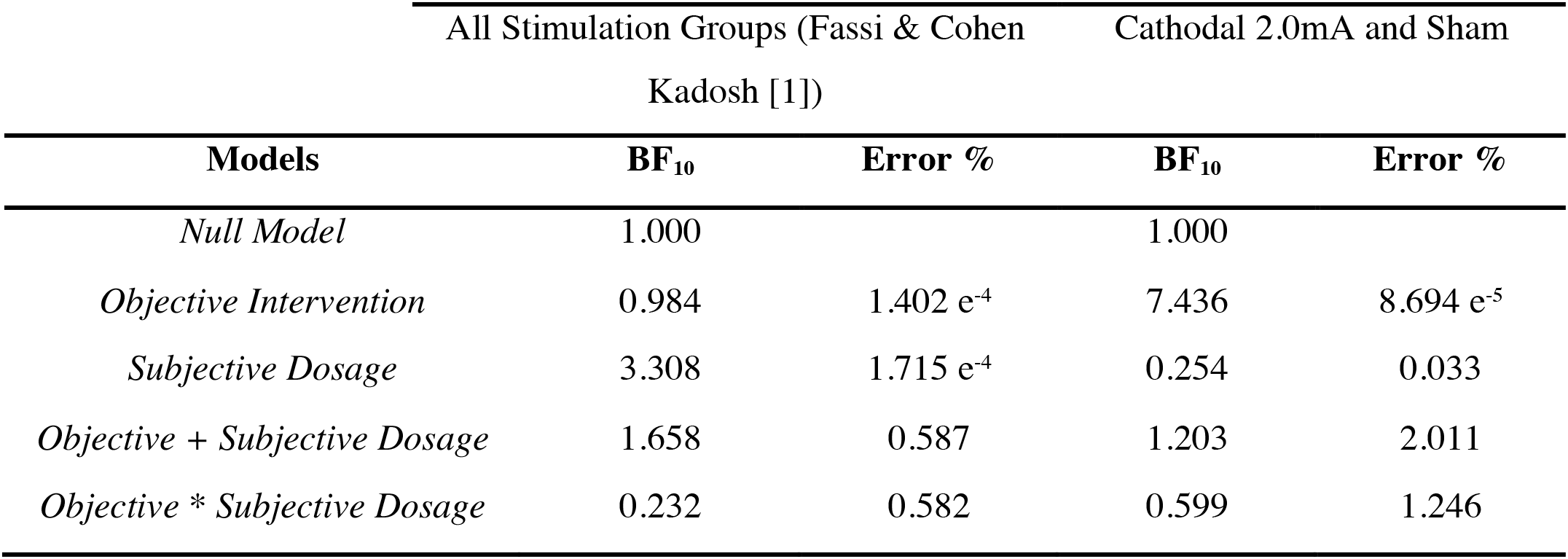
Comparison of the summarised Bayesian ANOVA of objective intervention and subjective dosage findings from Fassi & Cohen Kadosh [1] and the present study evaluating cathodal 2.0mA stimulation only.

### Comparing Non-Significant Conditions to Sham

After analysing each individual non-significant condition against sham in separate Bayesian ANOVAs comparing objective and subjective intervention, the results indicate that in the absence of an objective effect, objective intervention remained the ‘best’ model predictor for anodal 1.0mA (BF_10_= 0.781, *F*(1, 58)= 1.363, *p*=.248) and cathodal 1.5mA (BF_10_=2.189, *F*(1, 58)= 2.754, *p*=.103) compared to that of subjective intervention for the anodal 1.0mA (BF_10_= 0.458, *F*(1, 58)= 0.105, *p*=.572) or cathodal 1.5mA conditions(BF_10_= 0.788, *F*(1, 58)= 0.664, *p*= .419), although it should be noted that these BF values were anecdotal at best. Only within the cathodal 1.0mA condition did subjective intervention (BF_10_= 0.735, *F*(1, 58)= 1.310, *p*= .257) provide a higher BF value than objective intervention (BF_10_=0.540, *F*(1, 58)= 1.919, *p*= .171). However, given these BF values fall below 1 the null model is considered the best model predictor and these results are non-evidential. In the absence of an objective effect, subjective information has the potential to be a better predictor of results, and this should be considered by investigators. Even though this was not seen with our data, if a study is inadequately blinded then it is plausible that subjective experience *may* lead to significant differences in performance, and thus lead to spurious effects where no active control condition is included.

## Discussion

Fassi and Cohen Kadosh [1] tested the hypothesis that participants’ subjective beliefs about the influence of tDCS on a cognitive task drove the previous results of Filmer et al. [2] on mind-wandering. The authors concluded that subjective intervention, a participant’s personal belief about which stimulation condition they received, was a significant predictor of mind-wandering over and above that of objective intervention. Similarly, when investigating dosage, it was concluded that subjective dosage, a participant’s personal belief regarding the intensity of stimulation they received was also a better predictor of mindwandering than objective intervention alone. However, we argued that this was misleading given that Fassi and Cohen Kadosh [1] looked at all conditions from Filmer et al. [2] when only one stimulation condition, cathodal 2.0mA, produced meaningful effects regarding mind-wandering. In short, it was problematic to evaluate the effect of subjective belief across all stimulation conditions as this diluted the previous positive finding.

When evaluating *only* cathodal 2.0mA compared to sham from Filmer et al. [2] the effects of subjective beliefs about both intervention and dosage no longer are found. Specifically, we observed moderate evidence to support the notion that objective intervention is the strongest predictor of mind-wandering with higher rates of mind-wandering observed within the cathodal 2.0mA condition relative to sham. Interestingly, subjective intervention became the weakest predictor within the model, lesser than that of objective intervention, objective intervention and subjective intervention combined or an interaction of the two.

We found a similar pattern of results when evaluating dosage, as objective intervention was a better predictor of mind-wandering behaviour than subjective dosage or an interaction of the two. However, there is a caveat to this analysis. When comparing via an ANOVA, multiple levels of comparison are implemented to create multiple comparison conditions. In both the Fassi and Cohen Kadosh [1] paper as well as the present paper, there is an issue of group size in comparisons of overall stimulation conditions using dosage as there are far less participants who fall into the strong subjective dosage condition. For a full summary of group size in comparison conditions across all stimulation conditions in dosage see Appendix C1. This is further exacerbated when we compare only cathodal 2.0mA and sham stimulation as only one participant falls within the strong subjective dosage and sham objective dosage condition. The table in Appendix C2 outlines how this differs from other conditions, especially that of the sham objective/sham subjective condition, which contains 19 participants. This makes it difficult to draw strong conclusions from the results of the ANOVAs comparing dosage, as it is representative of only a single individual, not a sample of the population. Aspects such as these should be considered when choosing to apply these sorts of methods of evaluating subjective influences in the future to determine whether it is appropriate to pursue methods such as ANOVA. Given that it is only appropriate to further investigate conditions in which a significant change in behaviour has occurred (i.e., cathodal 2.0mA in this study), group size must be considered before further analysis.

Despite the present results, the concerns raised about subjective participant beliefs are warranted. Given the thought-provoking critique of the current standard methods of blinding within the literature, more sensitive measures and tests are needed to investigate issues of blinding and subjective belief. Indeed, further investigation of previous data sets and studies within the literature is needed to determine whether these findings have been influenced by participant belief.

Future studies should implement measures that aim to reduce the effects of subject belief and improve the blinding methods. For example, future studies should aim for the effective investigation and reporting of cutaneous perception, not just for safety, but also to gauge the perceptual experience of the participant and to elucidate sham efficacy. Investigators cannot draw conclusions about the efficacy of blinding, nor the subjective experiences of participants influencing behaviour, without first gaining insight into participant experience during experiments. There is also the possibility of implementing methods such as anaesthetic creams to remove the cutaneous feelings entirely from both active and sham conditions, a method that has been successfully used in both animal studies using transcranial alternating current stimulation (tACS) [27] a similar method to tDCS that involves an oscillating electrical current rather than a direct current [31] and in humans during tDCS [24] [28]. Another consideration is participant experience level, given the findings of Ambrus et al., [23] who observed that participants who had previous experience with tDCS were more likely to correctly identify trials which featured stimulation and those that featured sham whereas naïve participants were less likely to correctly identify their stimulation condition. This is a relatively easy to control measure through screening processes during recruitment that could help to reduce the unblinding of participants.

Relying on the comparison of a single target condition to a sham condition can be problematic if effective blinding is not achieved, particularly at the participant level as typically participants guess above chance in identifying stimulation conditions [2]. Further, the points raised by Fassi and Cohen Kadosh [1] and Filmer et al., [2] as discussed earlier, support the need for the departure from solely sham controlled methods. The use of active controls or a combination of both active and sham-controlled methods would avoid the issue of unblinding due to perceptual sensation. Active controls also allow for the investigation of the specific role of targeted brain regions when a separate brain region is targeted in the control, or polarity effects when an alternative polarity is applied to the control region [5]. Given the possibility to also investigate hemispheric differences, active controls can provide both a rich comparative measure and interesting investigative opportunities. However, when comparing two active conditions it can be difficult to distinguish which condition is modulating the behaviour when there is a difference between the two conditions, or whether both have an effect. Thus, the most beneficial configuration would be to include a condition of interest, a sham condition and at least one other active control condition, to help mitigate the pros and cons of each technique.

The insights and discussions outlined in the present study are facilitated by open science practices and underscore the importance and utility of sharing data. Without these practices, the field would continue to stagnate with poor methodological methods, particularly ineffective blinding, and the discussion of future improvements would be limited. Here we have made the results of Filmer et al. [2] more definitive, however this would not have been possible with the insights of Fassi and Cohen Kadosh [1] highlighting the benefits of multilab collaboration.

## Acknowledgements

This research was supported by an Australian Research Council (ARC) Discovery grant (DP210101977, PED & HLF). HLF was supported by an ARC Discovery Early Career Researcher Award (DE190100299). An Earmarked Research Training Program Stipend funded through The University of Queensland attached to an ARC Discovery Grant (DP210101977) supported MSG.

## Appendix A

**Appendix A1.** Bayesian ANOVA comparing Objective and Subjective Intervention for best model fit using average TUT score as the outcome measure [1].

| **Models** | **P(M)** | **P(M \| Data)** | **BF<sub>10</sub>** | **Error %** |
| --- | --- | --- | --- | --- |
| *Subjective Intervention* | 0.200 | 0.464 | 3.374 | 8.94 e<sup>-8</sup> |
| *Objective + Subjective Intervention* | 0.200 | 0.205 | 1.492 | 2.201 |
| *Null Model* | 0.200 | 0.137 | 1.000 |  |
| *Objective* | 0.200 | 0.135 | 0.984 | 1.40 e<sup>-4</sup> |
| *Objective \* Subjective Intervention* | 0.200 | 0.059 | 0.429 | 1.000 |

**Appendix A2.** Bayesian ANOVA comparing Objective and Subjective Dosage for best model fit using average TUT score as the outcome measure [1].

| **Models** | **P(M)** | **P(M \| Data)** | **BF<sub>10</sub>** | **Error %** |
| --- | --- | --- | --- | --- |
| *Subjective Dosage* | 0.200 | 0.489 | 3.308 | 1.715 e<sup>-4</sup> |
| *Objective + Subjective Dosage* | 0.200 | 0.219 | 1.658 | 0.587 |
| *Null Model* | 0.200 | 0.132 | 1.000 |  |
| *Objective* | 0.200 | 0.130 | 0.984 | 1.402 e<sup>-4</sup> |
| *Objective \* Subjective Dosage* | 0.200 | 0.031 | 0.232 | 0.582 |

## Appendix B

**Appendix B1.** Bayesian ANOVA comparing Objective and Subjective Intervention with the stimulation conditions Cathodal 2.0mA and Sham for best model fit using average TUT score as the outcome measure.

| **Models** | **P(M)** | **P(M \| Data)** | **BF<sub>10</sub>** | **Error %** |
| --- | --- | --- | --- | --- |
| *Null Model* | 0.200 | 0.376 | 1.000 |  |
| *Objective Intervention* | 0.200 | 0.639 | 7.436 | 8.694 e<sup>-5</sup> |
| *Objective Intervention + Subjective Intervention* | 0.200 | 0.183 | 2.133 | 2.627 |
| *Objective + Subjective Intervention + Objective Intervention \* Subjective Intervention* | 0.200 | 0.069 | 0.798 | 4.306 |
| *Subjective Intervention* | 0.200 | 0.024 | 0.276 | 0.004 |

**Appendix B2.** Bayesian ANOVA comparing Objective and Subjective Dosage with the stimulation conditions Cathodal 2.0mA and Sham for best model fit using average TUT score as the outcome measure.

| **Models** | **P(M)** | **P(M \| Data)** | **BF<sub>10</sub>** | **Error %** |
| --- | --- | --- | --- | --- |
| *Null Model* | 0.200 | 0.376 | 1.000 |  |
| *Objective Dosage* | 0.200 | 0.709 | 7.436 | 8.694 e<sup>-5</sup> |
| *Objective Dosage + Subjective Dosage* | 0.200 | 0.115 | 1.203 | 2.011 |
| *Objective + Subjective Dosage + Objective Dosage \* Subjective Dosage* | 0.200 | 0.057 | 0.599 | 1.246 |
| *Subjective Dosage* | 0.200 | 0.024 | 0.254 | 0.033 |

## Appendix C

**Appendix C1.** Frequency table for all stimulation conditions when comparing Objective Dosage and subjective dosage

| Subjective Dosage | Objective Dosage | Frequency |
| --- | --- | --- |
| Sham | Sham | 19 |
|  | Anode1mA | 8 |
|  | Cathode1mA | 11 |
|  | Cathode1.5mA | 7 |
|  | Cathode2.0mA | 12 |
|  | Missing | 0 |
|  | Total | 57 |
| Weak | Sham | 5 |
|  | Anode1mA | 6 |
|  | Cathode1mA | 8 |
|  | Cathode1.5mA | 5 |
|  | Cathode2.0mA | 5 |
|  | Missing | 0 |
|  | Total | 29 |
| Moderate | Sham | 5 |
|  | Anode1mA | 12 |
|  | Cathode1mA | 9 |
|  | Cathode1.5mA | 17 |
|  | Cathode2.0mA | 9 |
|  | Missing | 0 |
|  | Total | 52 |
| Strong | Sham | 1 |
|  | Anode1mA | 4 |
|  | Cathode1mA | 2 |
|  | Cathode1.5mA | 1 |
|  | Cathode2.0mA | 4 |
|  | Missing | 0 |
|  | Total | 12 |

**Appendix C2.** Frequency table for the stimulation conditions Cathodal 2.0mA and Sham (comparing subjective dosage and objective dosage)

Appendix C1. Frequency table for all stimulation conditions when comparing Objective Dosage and subjective dosage
| **Subjective Dosage** | **Objective Dosage** | **Frequency** |
| --- | --- | --- |
| Sham | Sham | 19 |
|  | Cathode2.0mA | 12 |
|  | Missing | 0 |
|  | *Total* | *31* |
| Weak | Sham | 5 |
|  | Cathode2.0mA | 5 |
|  | Missing | 0 |
|  | *Total* | *10* |
| Moderate | Sham | 5 |
|  | Cathode2.0mA | 9 |
|  | Missing | 0 |
|  | *Total* | *14* |
| Strong | Sham | 1 |
|  | Cathode2.0mA | 4 |
|  | Missing | 0 |
|  | *Total* | *5* |

## Notes

https://doi.org/10.14264/uql.2019.295

### Competing Interest Statement

The authors have declared no competing interest.

